# RNase 2/EDN cleaves tRNA anticodon loops to generate immunoactive RNAs

**DOI:** 10.1101/815803

**Authors:** Megumi Shigematsu, Atsushi Minami, Takuya Kawamura, Sushrut D. Shah, Tetsuhiro Ogawa, Deepak A. Deshpande, Yohei Kirino

**Affiliations:** Computational Medicine Center, Sidney Kimmel Medical College, Thomas Jefferson University, Philadelphia, Pennsylvania, 19107, USA; Department of Biochemistry and Molecular Biology, Sidney Kimmel Medical College, Thomas Jefferson University, Philadelphia, Pennsylvania, 19107, USA; Department of Biotechnology, The University of Tokyo, Tokyo, 113-8657, Japan; Center for Translational Medicine, Sidney Kimmel Medical College, Thomas Jefferson University, Philadelphia, Pennsylvania, 19107, USA; Collaborative Research Institute for Innovative Microbiology, The University of Tokyo, Tokyo, 113-8657, Japan

**Author notes:** Correspondence: Megumi Shigematsu, Ph.D., 1020 Locust Street, Philadelphia, PA 19107, USA; Yohei Kirino, Ph.D., 1020 Locust Street, Philadelphia, PA 19107, USA.

## Abstract

The RNase A superfamily, one of the most extensively studied enzyme families, remains incompletely defined with respect to its endogenous RNA substrates, recognition mechanisms, and physiological functions. Here, we identify eosinophil-derived neurotoxin (EDN/RNase 2) as a principal endoribonuclease mediating tRNA cleavage in the asthmatic lung. Inhalation of house dust mite (HDM) in mice induces robust accumulation of tRNA halves in the lung, coinciding with exclusive upregulation of EDN, but not other ribonucleases. In human lung epithelial cells, internalized EDN cleaves specific tRNAs and generates multiple species of tRNA halves, including immunostimulatory species that activate Toll-like receptor 7 (TLR7). EDN also promotes the release of extracellular vesicles enriched in these RNAs, implicating them in cytokine production during asthma pathogenesis. Biochemical analyses and structural simulations reveal that EDN recognizes the anticodon loop of tRNA by anchoring the phosphate backbones flanking the cleavage site, thereby precisely positioning the C34–A35 phosphodiester bond for catalysis. Furthermore, the highly efficient cleavage by EDN is mediated by conserved elements, including Arg36 in EDN and U33 and C38 in the target tRNA. Together, these findings establish the endogenous RNA substrate of EDN and uncover a molecular mechanism that links RNase A superfamily activity to immune response.

---

Asthma is characterized by irreversible structural and functional remodeling of the airways, driven by a complex network of inflammatory mediators, including cytokines, chemokines, and growth factors ^1–3^. Elucidating its molecular pathogenesis could provide improved diagnostic markers and therapeutic targets. While protein-based regulators such as transcription factors have been studied extensively for decades ^4,5^, short non-coding RNAs (sncRNAs), particularly tRNA halves—among the most abundant subclass of tRNA-derived sncRNAs—have only recently emerged as potent modulators of immune responses. Studies from our lab have shown that bacterial infection of human macrophages induces the production of tRNA halves and promotes the selective packaging of 5′-tRNA halves into extracellular vesicles (EVs) ^6,7^. Importantly, 5′-tRNA halves derived from tRNA^HisGUG^ and tRNA^ValCAC/AAC^ (5′-HisGUG and 5′-ValCAC/AAC) potently activate Toll-like receptor 7 (TLR7) upon delivery into recipient cells, thereby triggering cytokine production ^6–9^. These findings suggest that tRNA halves function as endogenous immunostimulatory ligands capable of shaping immune response.

The accumulation of tRNA half molecules has been observed across diverse cell types, tissues, and biofluids in various physiological and pathological contexts ^10–12^, suggesting broad biological relevance and a regulated biogenesis mediated by specific endoribonucleases that cleave the tRNA anticodon loop. However, the endogenous functions of these endoribonucleases and the molecular basis of their substrate recognition remain poorly understood. Although immunostimulatory tRNA halves have been implicated in infectious and inflammatory diseases such as *Mycobacterium tuberculosis* infection ^8^ and chronic obstructive pulmonary disease (COPD) ^9^, their enzymatic origins remain largely obscure. The inability of conventional RNA-seq to capture 5′-tRNA halves bearing a 2′,3′-cyclic phosphate (cP), a hallmark of cleavage by anticodon loop-targeting endonucleases such as Angiogenin ^13–15^, RNase Kappa ^16^, RNase L ^17,18^, and IRE1α ^19–21^, has contributed not only to the scarcity of tRNA half profiling data but also to the limited characterization of their biogenesis-associated endoribonucleases.

## tRNA halves are enriched in asthmatic lungs

To characterize tRNA halves in asthma, we employed a widely used house dust mite (HDM) inhalation model in mice ^22^. HDM exposure induced airway remodeling, including upregulation of α-smooth muscle actin (α-SMA) (**Extended Data Fig. 1a**), confirming the establishment of an asthmatic phenotype ^23^. Total RNA extracted from the lungs of HDM-challenged mice showed a prominent accumulation of sncRNAs approximately 30–40 nucleotides (nt) in length (**Fig. 1a**), and northern blot showed the increased accumulation of tRNA halves (**Fig. 1b**). For tRNA half profiling, we applied cP-RNA-seq, a sequencing method optimized to capture sncRNAs bearing cP termini, such as 5′-tRNA halves ^14,24,25^ (**Extended Data Fig. 1b**). Analysis of tRNA-derived reads from the cP-RNA-seq libraries of HDM-treated lungs revealed a bimodal length distribution, with peaks at 34 and 38 nt corresponding to 5′- and 3′-tRNA halves, respectively (**Fig. 1c**). Subclassification into nine tRNA fragment (tRF) subtypes (**Fig. 1d**) identified 5′-tRNA halves as the most abundant species, followed by 3′-tRNA halves (**Fig. 1e**). Note that cP-RNA-seq does not detect aminoacylated 3′-tRNA halves; instead, it primarily detects 3′-tRNA halves or 3′-tRFs lacking the 3′-terminal adenine at nucleotide position (np) 76 (A76) (**Extended Data Fig. 1c**), as previously reported ^26,27^.

**Figure 1.**
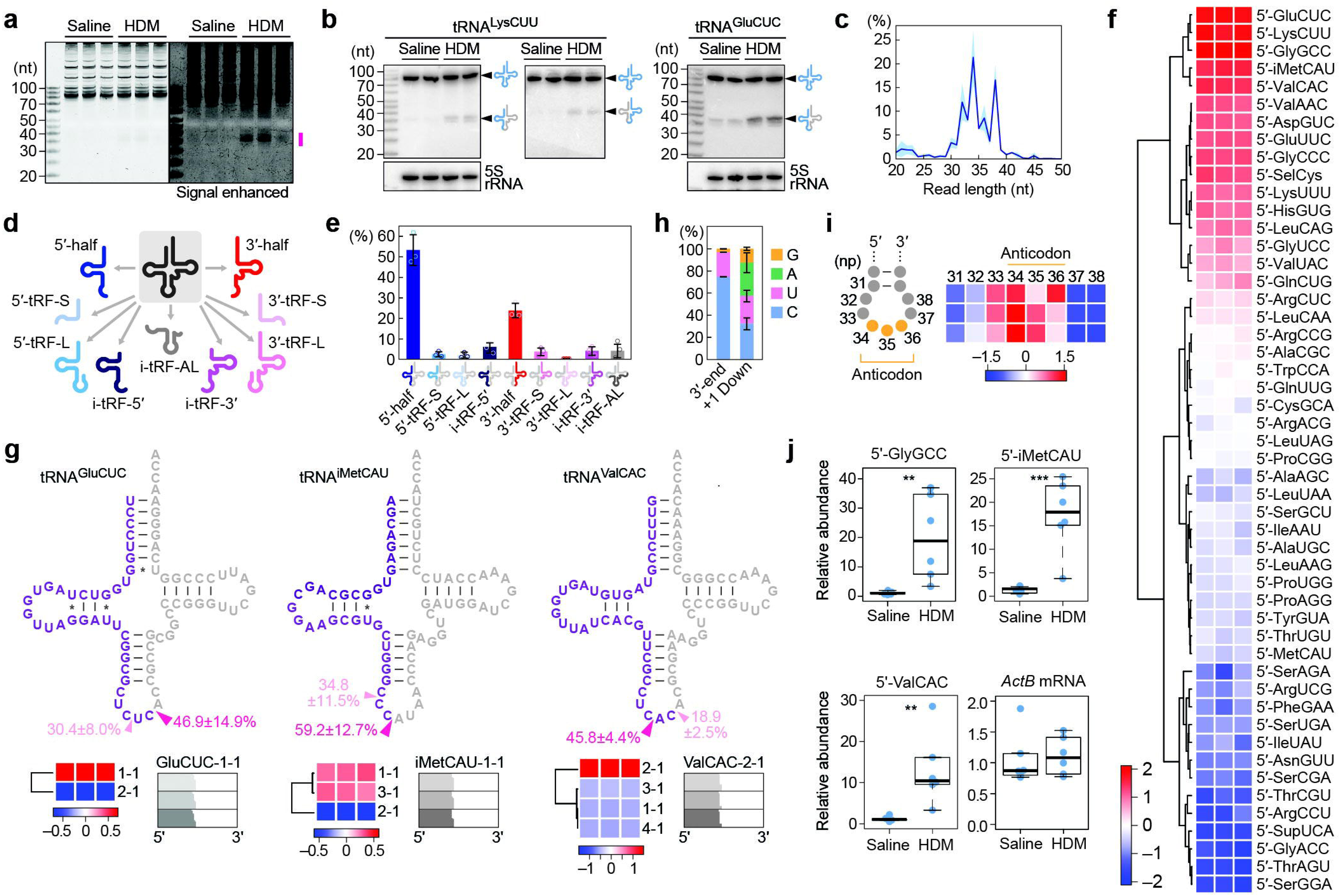
tRNA halves accumulate in the lungs of HDM-challenged mice. **(a)** Total RNA from mouse lungs was stained with SYBR Gold. The region enriched in sncRNAs in HDM-challenged is indicated (magenta line). **(b)** Northern blot detecting 5′-LysCUU, 3′-LysCUU, and 5′-GluCUC (left to right). **(c)** Length distribution of tRNA-mapped reads from cP-RNA-seq of HDM-challenged lungs. Shaded area indicates standard deviation (S.D.) across three biological replicates. **(d)** Schematic of nine tRNA fragment subclasses. 5′- and 3′-halves retain the 5′- and 3′-ends of mature tRNAs, respectively, and result from anticodon loop cleavage. 5′- and 3′-tRFs also retain the mature tRNA ends but are generated from cleavage outside the anticodon loop, either excluding (short, S) or including (long, L) the anticodon loop. Internal fragments (i-tRFs) lack mature ends, derived from the 5′-(i-tRF-5′) and 3′-half (i-tRF-3′) regions or the region encompassing the anticodon loop (i-tRF-AL). **(e)** Proportion of tRNA-mapped reads by subclass. Error bars, S.D. from three replicates. **(f)** Heatmap of 5′-tRNA half abundance across isoacceptors. Color bar indicates z-scores. **(g)** Cleavage patterns for 5′-tRNA halves from selected isoacceptors. Cloverleaf structures indicate major cleavage sites (arrowheads). Errors represent the S.D. from three replicates. Heatmaps show relative abundance of isodecoders; most abundant isodecoder alignments are also shown. **(h)** Nucleotide composition at the 3′-end and the immediately downstream position of 5′-tRNA halves. **(i)** Frequency of 3′-end positions of 5′-tRNA halves. Nucleotide position numbering follows Sprinzl’s nomenclature. Color scale represents z-scores. **(j)** Quantification of select 5′-tRNA halves by TaqMan qPCR, normalized to U6 snRNA. ActB mRNA was quantified by standard qPCR and also normalized to U6 snRNA. Sample number: n=6 for each cohort (one saline sample for 5′-ValCAC was not detected). Student’s *t*-test: \*\**p*<0.01; \*\*\**p*<0.001.

The accumulation of 5′-tRNA halves in HDM-treated lungs was largely attributable to a restricted subset of species, rather than being evenly derived from all 51 tRNA isoacceptors. Twelve dominant species accounted for over 90% of all 5′-tRNA halves detected (**Fig. 1f**), among which 5′-ValCAC, 5′-ValAAC, and 5′-HisGUG are known immunostimulatory RNAs capable of activating TLR7 ^6–9^. Isodecoder-level analysis confirmed that 5′-tRNA halves originated from a defined subset of tRNA genes (**Fig. 1g and Extended Data Fig. 1d**). Although multiple sites were cleaved within individual tRNA species, the predominant site for 5′-tRNA half production was the 3′-side of the cytosine (C), followed by uracil (U), indicating a strong pyrimidine preference (**Fig. 1h**). The downstream nucleotide was most often adenine (A), followed by C and U, suggesting preferential cleavage between specific nucleotide pairs such as C–A (**Fig. 1h**). The 3′-ends of 5′-tRNA halves were largely confined to the anticodon region, with strong enrichment at np 34 (the anticodon first position), whereas cleavage was infrequent at np 37 and 38 (**Fig. 1i**).

We further confirmed the upregulation of selected tRNA half species using TaqMan qPCR, which enables specific quantification of tRNA halves ^14,28,29^. This analysis revealed marked increases in the levels of all examined 5′-tRNA halves, 5′-GlyGCC, 5′-iMetCAU, and 5′-ValCAC, in HDM-exposed lungs (**Fig. 1j**). Together, these results demonstrate an abundant accumulation of specific tRNA halves in the lungs of HDM-induced asthmatic mice, driven by site-selective cleavage within tRNA anticodon loops.

## *EDN* is specifically upregulated in asthma

To identify the enzyme responsible for tRNA half generation, we systematically profiled ribonuclease gene expression in lungs from HDM-challenged mice. We compiled 49 mRNA-seq datasets from four independent studies ^30–32^ and performed differential expression analyses, focusing on 126 genes annotated as ribonucleases. This analysis revealed a striking and consistent upregulation of *Ear11* (*<u>E</u>osinophil-<u>a</u>ssociated <u>r</u>ibonuclease 11; RNase 2a*), the rodent ortholog of human *<u>e</u>osinophil-<u>d</u>erived <u>n</u>eurotoxin* (*EDN, RNASE2*), in HDM-treated lungs across all four studies (**Fig. 2a**).

**Figure 2.**
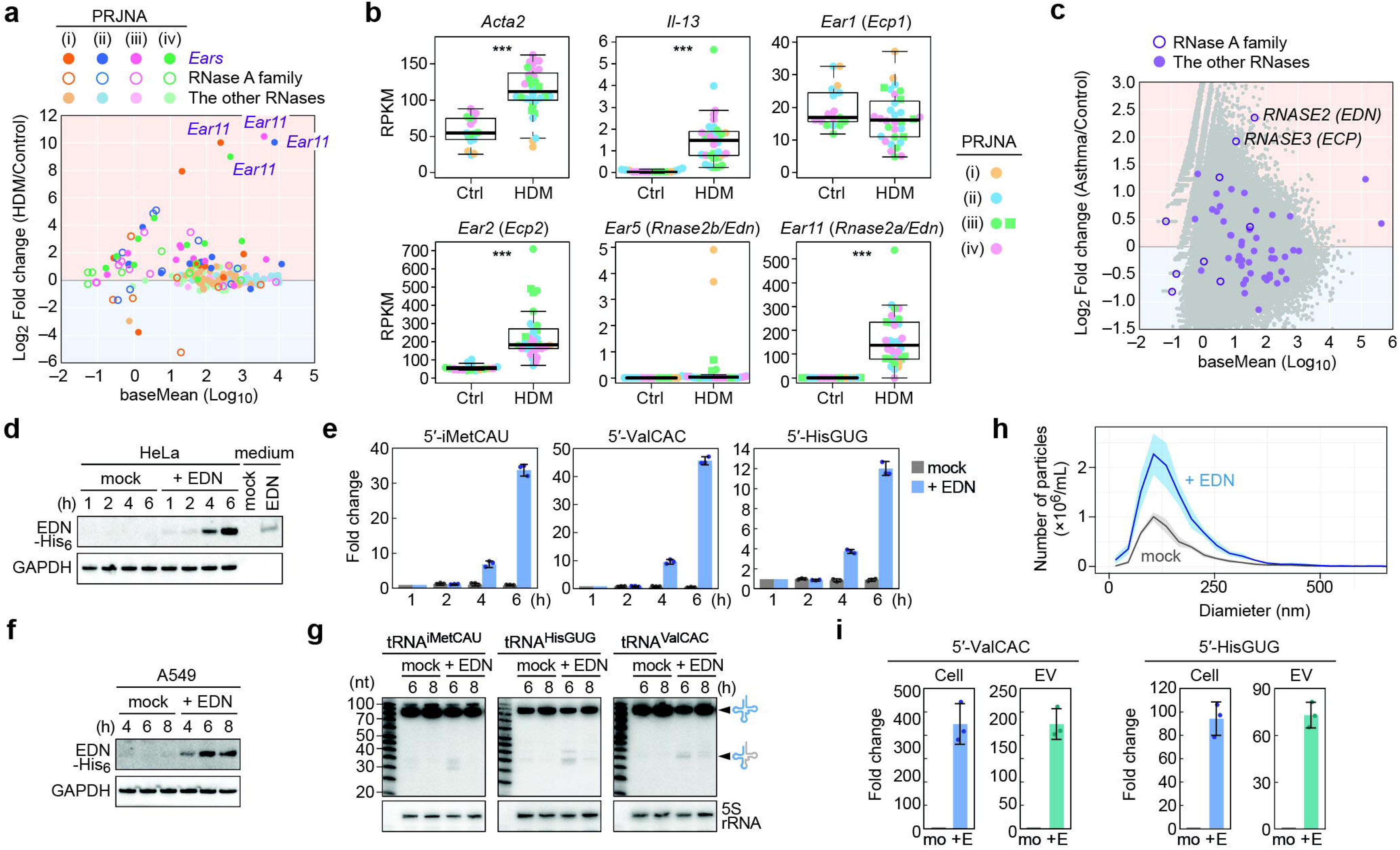
EDN produces tRNA halves in lung epithelial cells. **(a)** MA plot of mouse genes showing the differential expression in the lungs of HDM-challenged versus control mice. Genes annotated as ribonucleases were extracted and shown. The x-axis indicates the mean expression level of each gene, and the y-axis represents fold change. Four independent datasets were analyzed: (i) PRJNA212775 ^30^; (ii) PRJNA547063 ^31^; (iii) PRJNA522698 ^31^; and (iv) PRJNA480707 ^32^. **(b)** mRNA expression levels (RPKM) of *Acta2* and *Il-13* (HDM-positive controls) and five *Ears* annotated as *Edn (RNase 2)* or *Ecp*. Color coding corresponds to panel (a). In PRJNA480707, low and high HDM does are shown as green squares and circles, respectively. Student’s *t*-test: \*\*\**p*<0.001. **(c)** MA plot of ribonuclease gene expression in peripheral blood of asthma patients versus healthy controls, obtained by analyzing dataset of PRJNA415959 ^35^. **(d)** Western blot analysis of recombinant EDN uptake by HeLa cells over time. **(e)** TaqMan qPCR quantification of 5′-iMetCAU and 5′-HisGUG in HeLa cells treated as in (d), normalized to 5S rRNA. Error bars, S.D. from three technical replicates of qPCR. **(f)** Western blot of recombinant EDN uptake by A549 cells over time. **(g)** Northern blot detection of 5′-tRNA halves in A549 cells treated as in (f). **(h)** NTA of EVs secreted from A549 cells after 8 h of EDN treatment. Shaded region represents S.D. across three independent experiments. **(i)** TaqMan qPCR quantification of 5′-ValCAC and 5′-HisGUG in A549 cells and secreted EVs, normalized to 5S rRNA (cells) or to equal isolated volume (EVs). Error bars, S.D. from three independent experiments.

*EDN* belongs to the RNase A superfamily, which in humans comprises 13 members, each encoded by a single gene ^33^. In contrast, rodents possess expanded *Ear* genes ^34^ (**Extended Data Fig. 2a**), functionally analogous to human *EDN* and *<u>e</u>osinophil <u>c</u>ationic <u>p</u>rotein* (*ECP, RNASE3*). Given the lack of systematic expression profiling of *Ear* or RNase A superfamily genes in mouse tissues, and the lack of antibodies capable of distinguishing individual Ear proteins, we analyzed their transcript levels individually. Among them, *Ear11* and *Ear2* (*Ecp2*, the rodent ortholog of human *ECP*) showed the prominent basal expression and strongest induction in HDM-challenged lungs, with more than 500-fold and 3.98-fold upregulation, respectively (**Fig. 2b and Extended Data Fig. 2b**). Several other genes showed modest induction; however, their overall expression levels remained low.

We also analyzed clinical transcriptomic datasets from patients with asthma following exacerbation ^35^. Among 55 ribonucleases identified in differential expression analysis between healthy controls and asthmatics, *EDN* and *ECP* emerged as the most and second most upregulated genes in patients with asthma, respectively, while others showed minimal change or low expression (**Fig. 2c**). Both *EDN* and *ECP* are exclusively expressed in eosinophils, whose blood counts are elevated in asthma ^36,37^. Accordingly, serum levels of EDN and ECP are elevated in patients with asthma and show a positive correlation with disease severity ^38–41^. Given that EDN exhibits approximately 50-fold higher endoribonucleolytic activity than ECP ^42,43^, and that *Ear11 (EDN)* is more strongly induced than *Ear2 (ECP)* in HDM-treated lungs (**Fig. 2b**), we conclude that EDN is the primary endoribonuclease responsible for cleaving selective tRNA species and driving the accumulation of tRNA halves in asthmatic lungs.

## EDN cleaves tRNAs to generate and secrete immunoactive tRNA halves

Human EDN is a 15-kDa endoribonuclease comprising 134 amino acids and harboring three conserved catalytic residues (His15, Lys38, and His129) essential for its enzymatic activity ^44^. A previous study reported that EDN can cleave tRNAs ^45^; however, its *in vivo* relevance and substrate specificity remain unclear. To evaluate its intracellular activity, we purified recombinant human EDN expressed in *E. coli* using a denaturing protocol followed by stepwise refolding (**Extended Data Fig. 3a**). Incubation of the recombinant EDN with HeLa total RNA resulted in substantial RNA degradation, confirming the ribonucleolytic activity of EDN *in vitro* (**Extended Data Fig. 3b and 3c**). Treatment of HeLa cells with recombinant EDN revealed a time-dependent intracellular accumulation, with markedly higher levels at 6 hours than at 4 hours, as assessed by western blot (**Fig. 2d, Extended Data Fig. 3d**). This increase in EDN paralleled the induction of 5′-tRNA halves, with significantly higher levels detected at 6 hours compared to 4 hours, as measured by TaqMan qPCR (**Fig. 2e**). Similar uptake and accumulation dynamics were observed in A549 lung epithelial cells, with peak EDN levels at 6 hours (**Fig. 2f**). Northern blot confirmed robust induction of all examined 5′-tRNA halves, with peak accumulation coinciding with intracellular EDN levels (**Fig. 2g**). Notably, two immunoactive tRNA halves, 5′-HisGUG and 5′-ValCAC, were abundantly generated by EDN in A549 cells.

While EDN is known to be released from eosinophils into the airways and to activate dendritic cells ^46,47^ , the role of EDN in tRNA cleavage and its relevance to asthma pathogenesis has remained undefined. Given the central role of lung epithelial cells in immune surveillance ^48,49^ and EV secretion ^50,51^, we investigated whether EDN influences EV biogenesis in A549 cells. Upon EDN treatment, nanoparticle tracking analysis (NTA) revealed a significant increase in EV release (**Fig. 2h, Extended Data Fig. 3e**), accompanied by a drastic enrichment of immunoactive 5′-HisGUG and 5′-ValCAC not only intracellularly but also within secreted EVs (**Fig. 2i**). These findings suggest that EDN not only generates immunoactive tRNA halves but also promotes EV biogenesis and facilitates the loading of these RNAs into EVs from lung epithelial cells, potentially contributing to the inflammatory process in asthma.

## EDN selectively generates specific tRNA half species

Because the mouse asthma model involves prolonged HDM treatment, the observed tRNA half profiles likely reflect not only tRNA cleavage but also post-cleavage dynamics such as RNA stability and turnover. To determine the primary cleavage events mediated by EDN, which leaves cP at the cleavage site ^42^, we treated BEAS-2B human lung epithelial cells with recombinant EDN (**Fig. 3a**) and profiled the resulting sncRNAs using cP-RNA-seq. In EDN-treated cells, 5′-tRNA halves represented the most abundant subclass among tRNA-derived reads, followed by 3′-tRNA halves (**Fig. 3b**), recapitulating the patterns seen in HDM-exposed mouse lungs (**Fig. 1e**). Comparative analysis with mock-treated controls revealed that a specific subset of 5′-tRNA halves was preferentially enriched upon EDN treatment. Marked increases were observed for 5′-GluCUC/UUC, 5′-ValCAC, 5′-iMetCAU, 5′-HisGUG, 5′-AspGUC, and 5′-LeuCAG, suggesting that their parent tRNAs are favored substrates of EDN (**Fig. 3c and Extended Data Fig. 4**). In contrast, typically abundant species such as 5′-LysCUU and 5′-GlyGCC were relatively downregulated, indicating lower production efficiency by EDN. Notably, these observations reflect relative enrichment within libraries rather than absolute RNA abundance; accordingly, even tRNA halves derived from less-favored substrates exhibited increased absolute levels following EDN treatment. Mapping of cleavage sites revealed that EDN-enriched 5′-tRNA halves were predominantly generated by cleavage at np 34 within the anticodon loop, a pattern distinct from that of downregulated species (**Fig. 3d**). The immunoactive 5′-ValCAC and 5′-HisGUG were primarily generated by cleavage at C34–A35 and U33–G34, respectively, with additional major cleavage observed at C36–A37 and U35–G36 (**Fig. 3e**), consistent with *in vivo* pattern observed in HDM-challenged mice (**Fig. 1g and Extended Data Fig. 1d**). These results indicate that cellular EDN activity preferentially cleaves at or near np 34 within the anticodon loop, with substrate specificity shaped by the sequence and structure of individual tRNAs.

**Figure 3.**
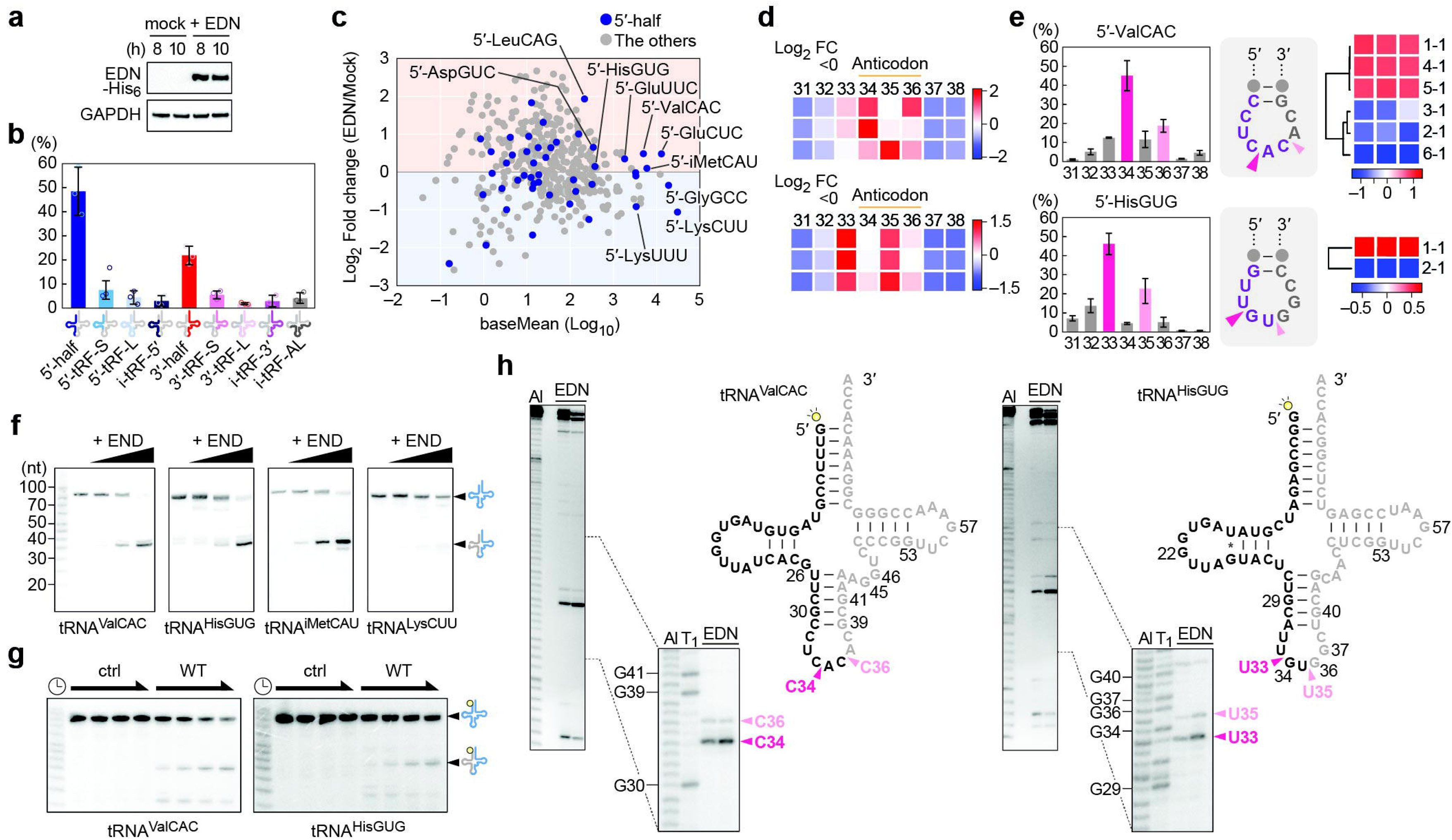
Characterization of EDN-generated tRNA halves. **(a)** Western blot showing uptake of recombinant EDN by BEAS-2B cells. **(b)** Classification of tRNA-mapped reads from EDN-treated BEAS-2B cells into nine subclasses. Error bars, S.D. from three biological replicates. **(c)** MA plot of 5′-tRNA halves (blue) and other subclasses of tRNA-derived sncRNAs (gray) aggregated by isoacceptors, comparing EDN-treated and mock-treated BEAS-2B cells. **(d)** Frequency of 3′-end positions of 5′-tRNA halves upregulated (top) or downregulated (bottom) upon EDN treatment. Color scale represents z-scores. **(e)** Bar graphs and ASL secondary structure showing proportion of 3′-end positions of 5′-tRNA halves. Heatmaps show the relative abundance of isodecoders. **(f)** Northern blot detecting 5′-tRNA halves generated by recombinant EDN cleavage of HeLa tRNA fractions. Reactions were performed at pH 5.5 for 1 min with 0, 1, 5, and 25 nM of EDN and 500 ng of tRNA fractions. **(g)** Recombinant EDN was incubated with 5′-^32^P-labaled mature tRNA^ValCAC^ and tRNA^HisGUG^, and the resulting 5′-tRNA halves were observed. Reactions were performed at pH 6.0 with 0.5 nM EDN and 250 nM substrate for 15, 30, 60, and 90 s. **(h)** The 30- and 90-s samples from (g) were analyzed by high-resolution PAGE. Al, alkaline ladder; T1, RNase T1 digested. Cleavage sites are indicated on each cloverleaf structure.

## Anticodon loops are the primary sites of EDN-mediated cleavage in tRNAs

To further investigate EDN’s substrate specificity, we reconstituted *in vitro* tRNA cleavage reactions using recombinant EDN. Incubation of gel-purified HeLa tRNAs with EDN, followed by northern blot, revealed the efficient generation of 5′-ValCAC, 5′-HisGUG, and 5′-iMetCAU, whereas 5′-LysCUU was produced to a much lesser extent (**Fig. 3f**). This species-selective cleavage preference was consistent with the profiles observed in EDN-treated BEAS-2B cells (**Fig. 3c**). EDN exhibited increased ribonucleolytic activity under mildly acidic conditions, peaking at pH 6.0 (**Extended Data Fig. 5a and 5b**), and thus a slightly acidic pH was used for further biochemical characterization.

*In vitro* EDN-mediated cleavage of synthetic 5′-^32^P-labeled tRNA^ValCAC^ and tRNA^HisGUG^ produced prominent tRNA halves (**Fig. 3g**). High-resolution denaturing PAGE mapped the primary cleavage site of tRNA^ValCAC^ at C34–A35, with a secondary site at C36–A37. For tRNA^HisGUG^, the major cleavage occurred at U33–G34, followed by U35–G36 (**Fig. 3h**). These cleavage patterns closely mirrored those identified by cP-RNA-seq in HDM-challenged mouse lungs and EDN-treated BEAS-2B cells (**Fig. 1g and 3e**), validating cP-RNA-seq as a reliable method for capturing primary EDN cleavage sites. Because synthetic tRNAs lack post-transcriptional modifications and may not fully adopt the native L-shaped tertiary structure, these results suggest that EDN does not require intact tRNA architecture for substrate recognition. Instead, EDN appears to recognize local features of the anticodon loop and preferentially cleaves at defined positions within this region.

## EDN preferentially cleaves between C and A on the 5′-side of the anticodon loop

To dissect the substrate preference of EDN at the anticodon loop, we employed DNA– RNA chimeric anticodon stem-loop (ASL) constructs based on the tRNA^ValCAC^ sequence, with a single RNA nucleotide at position 10 corresponding to tRNA np 34. Wild-type EDN efficiently cleaved the ASL substrate, whereas its catalytically inactive K38R mutant did not (**Fig. 4a**), confirming that the observed cleavage is attributable to EDN’s intrinsic catalytic activity. Among the 12 dinucleotide combinations tested at the cleavage site, EDN cleaved exclusively after pyrimidines, with rC–A showing the highest sensitivity, followed by rU–A (**Fig. 4b-c**). This pyrimidine preference at the 5′-side of the cleavage site is believed to be a conserved feature of RNase A superfamily members ^42,43^, and is likewise observed in EDN. The ranking of the four most efficiently cleaved dinucleotides mirrors that of RNase A but differs from Angiogenin and ECP ^42,52,53,54^, suggesting a conserved substrate recognition mechanism more closely aligned with RNase A than with the other members.

**Figure 4.**
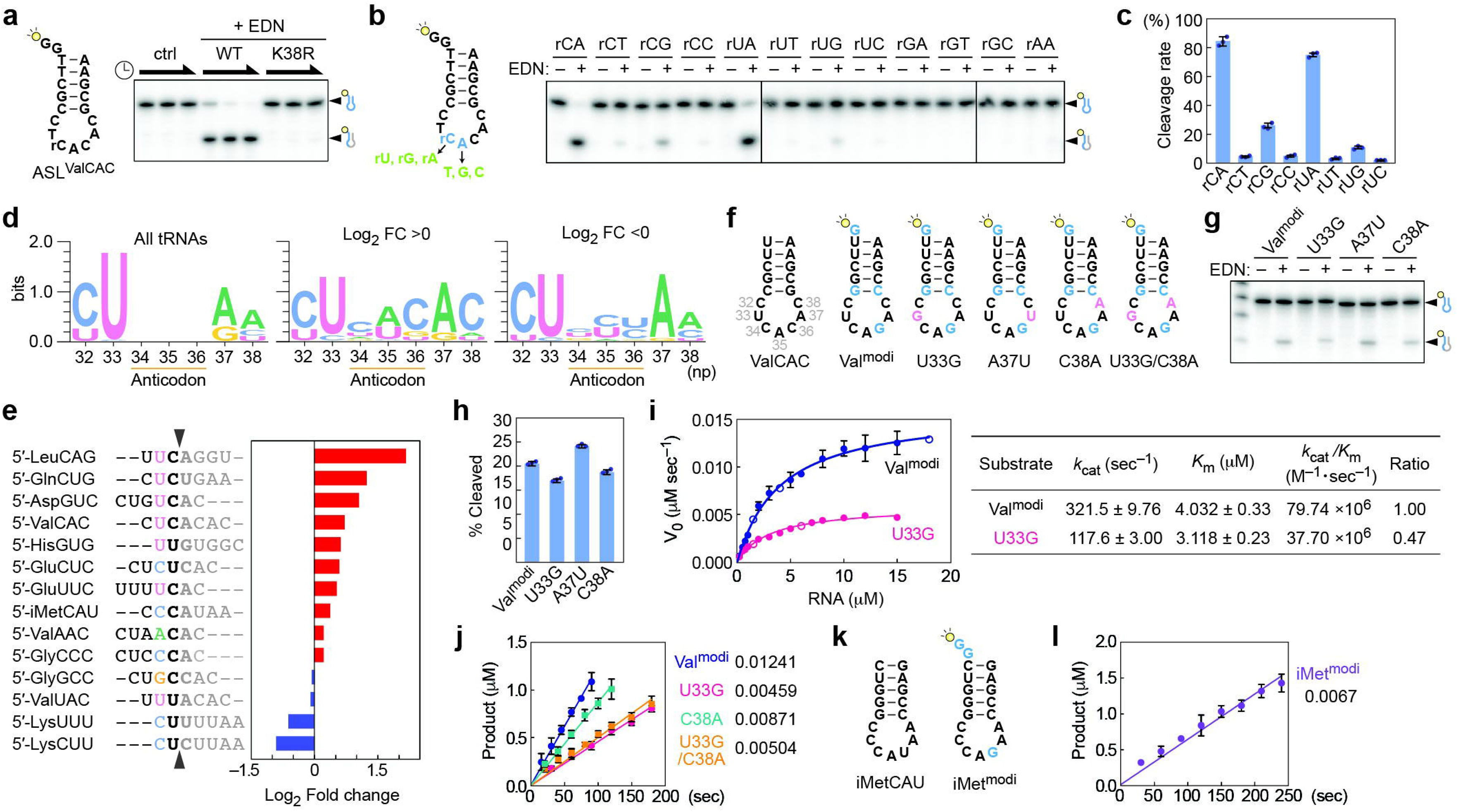
Enzymatic characterization of EDN cleavage of the tRNA anticodon loop. **(a)** Secondary structure of a DNA–RNA chimeric ASL^ValCAC^ (left) and its EDN cleavage products detected by PAGE (right). The ASL contained a single RNA residue (rC) at np 34 to define the cleavage site at rC34–A35. Reactions were performed either wild-type (WT) or K38R mutant EDN (0.5 nM) and 250 nM substrate for 10, 30, and 60 s. **(b)** ASL^ValCAC^ variants with defined dinucleotides at the cleavage site (left), and corresponding EDN-mediated *in vitro* cleavage results (right). **(c)** Quantification of cleavage efficiency for the ASL variants in (b). Error bars, S.D. from three independent experiments. **(d)** Nucleotide composition at np 32–38 of the anticodon loop in source tRNAs of BEAS-2B 5′-tRNA halves identified by cP-RNA-seq. Frequencies are shown for all 193 detected isodecoders (left), and for the 17 upregulated (> 100 RPM, middle) and 21 downregulated (>100 RPM, right) species upon EDN treatment. **(e)** Anticodon loop sequences and cleavage sites for 5′-tRNA halves, ranked by EDN cleavage efficiency as determined from BEAS-2B cP-RNA-seq data. **(f)** Secondary structure of ASL^ValCAC^ (all RNA) and its variants used in *in vitro* EDN cleavage assays. Nucleotide substitutions for detection purposes are shown in light blue, whereas mutations introduced to investigate their roles in cleavage are shown in pink. **(g)** *In vitro* cleavage of the indicated ASL constructs (2 μM) by EDN (100 pM, reaction time: 1 min). **(h)** Quantification of cleavage efficiency in (g). Error bars, S.D. from three independent experiments. **(i)** Michaelis-Menten plots for Val^modi^ and U33G ASL substrates (left) and the corresponding kinetic parameters (right). **(j)** Initial reaction rates of ASL variants (10 μM). **(k)** Secondary structure of ASL^iMetCAU^ (all RNA) and its variant used in the *in vitro* cleavage assay. **(l)** Initial reaction rates of iMet^modi^ (10 μM).

We next systematically shifted the RNA nucleotide position within the tRNA^ValCAC^-based ASL constructs to map positional preferences across the loop. Cleavage was most efficient at rC33–A34, followed by rC32–A33 and rC34–A35 (**Extended Data Fig. 5c and 5d**). Cleavage efficiency declined on the 3′-side of the loop, including rC36–A37 and rC37–A38. Similar patterns were observed in ASLs derived from tRNA^HisGUG^, where rU33–G34 was the most efficiently cleaved, followed by rU32–G33 and rU34–G35, with minimal cleavage at rU37–G38 (**Extended Data Fig. 5e and 5f**). These results suggest that EDN preferentially targets the 5′-side of the anticodon loop, implicating a structural determinant in substrate recognition by EDN.

Analysis of dinucleotide composition in human tRNAs revealed a relatively even distribution at np 34–35, except for underrepresented combination such as A–A, A–G, G–A, and G–G (**Extended Data Fig. 5**g). In contrast, np 36–37 exhibited strong compositional bias, with a significant enrichment of C–A and U–A (**Extended Data Fig. 5g**), both among the most preferred EDN substrates. If sequence alone dictated EDN cleavage, these 3′-side positions would be expected to serve as major targets. However, biochemical and cP-RNA-seq analyses demonstrated preferential cleavage at the 5′-side of the anticodon loop. For example, in tRNA^ValCAC^, EDN preferentially cleaves at C34–A35 over C36–A37. These findings suggest that cleavage specificity is not solely determined by dinucleotide composition but also critically influenced by local RNA structure and positional context within the anticodon loop.

## EDN recognizes a conserved nucleotide motif within the anticodon loop

tRNAs are among the most conserved RNAs, with certain nucleotides preserved across isoacceptors. In human tRNAs, U33 is highly conserved, and C32 shows modest conservation (**Fig. 4d**). tRNAs preferentially cleaved by EDN, identified via cP-RNA-seq (**Fig. 3c**), showed a notable enrichment of C38, in place of the more typical A38 (**Fig. 4d**). Furthermore, 5′-tRNA haves with a U residue immediately upstream of the cleavage site tended to exhibit higher fold increases upon EDN treatment (**Fig. 4e**). These findings suggest that EDN recognizes its substrates not only through the dinucleotide at the cleavage site, but also via the upstream and downstream sequence context. To test this, we utilized a series of all-RNA ASL constructs based on tRNA^ValCAC^. To enforce stable stem-loop formation and specific cleavage at C34–A35, we designed a reference construct Val^modi^ incorporating three modifications: addition of a G at the 5′-terminus, a C36G substitution, and inversion of a C–G base pair within the anticodon (**Fig. 4f**). Mutation of U33 to G (U33G) or C38 to A (C38A) impaired EDN cleavage, suggesting that these residues contribute to substrate recognition, whereas mutation of A37 to U (A37U) did not significantly affect cleavage efficiency (**Fig. 4f-h**).

Kinetic analyses revealed that Val^modi^ is a highly efficient EDN substrate, with a *k*_cat_ of 321.5 sec^−1^ and a *K*_m_ of 4.03 μM (**Fig. 4i**). This *K*_m_ is comparable to the cellular concentration of tRNAs, indicating that EDN can function at or near its maximal catalytic rate *in vivo*. These parameters are superior compared to those reported for EDN, ECP, and RNase A using dinucleotides substrates or yeast-derived tRNA pools ^44,54,55^, indicating that structured ASL constructs provide a more favorable context for ribonuclease activity. The U33G mutant exhibited a reduced *k*_cat_ of 117.6 sec^−1^, with little change in *K*_m_, resulting in a ∼53% decrease in catalytic efficiency (**Fig. 4i**). While this suggests that U33 enhances cleavage, the lack of *K*_m_ change implies that U33 may not serve as a direct contact point, but instead influence local structure and loop dynamics. The C38A mutation modestly reduced the cleavage rate (**Fig. 4j**), likely due to the formation of weaker hydrogen bonds between A38 and the conserved C32, which is known to lead to a more compact anticodon loop structure ^56^. The double mutant U33G/C38A did not show additive impairment beyond U33G alone (**Fig. 4j**), further highlighting the dominant role of U33 in regulating cleavage.

Notably, in cP-RNA-seq data from EDN-treated BEAS-2B cells, 5′-iMetCAU exhibited only moderate upregulation compared to other efficiently generated 5′-tRNA halves (**Fig. 3c**), despite harboring the C34–A35 cleavage motif. The presence of C33, rather than the more conserved U33, in tRNA^iMetCAU^ may underlie its reduced susceptibility to EDN cleavage. To examine this, we constructed a modified ASL based on tRNA^iMetCAU^ by introducing a G36 mutation to match the Val^modi^ configuration (**Fig. 4k**). This iMet^modi^ construct, equivalent in loop sequence to Val^modi^ C38A but containing C33, showed even lower cleavage efficiency than the C38A mutant of Val^modi^ (**Fig. 4l**), providing further evidence for the functional importance of U33 in facilitating EDN-mediated cleavage.

## EDN forms a stable complex with anticodon loop through its catalytic and substrate-recognition residues

The rapid turnover of ribonucleases poses a challenge for experimentally capturing transient enzyme–substrate interactions. To elucidate how EDN recognizes and cleaves the anticodon loop, we performed molecular docking and molecular dynamics (MD) simulations. As an initial model, we docked the ASL of tRNA^ValCAC^, hereafter referred to as ASL^ValCAC^, into the active site of EDN, guided by the structure of *Thermus thermophilus* tRNA^ValCAC^ (PDB 1IVS) ^57^ and RNase A bound to d(CpA) (PDB 1RPG) ^58^. An AlphaFold3 (AF3) ^59^-predicted EDN–ASL complex model (**Extended Data Fig. 6a**) yielded a similar configuration, supporting the plausibility of the docked model. During 100-ns MD simulations, the overall orientation of ASL relative to EDN remained stable, with the stem-loop stably positioned across the catalytic groove (**Fig. 5a and 5b**). The low overall root mean square deviation (RMSD) values (<4) (**Extended Data** Fig. 6b) and favorable binding free energy (**Supplementary Table S1**) indicated a stable interaction.

**Figure 5.**
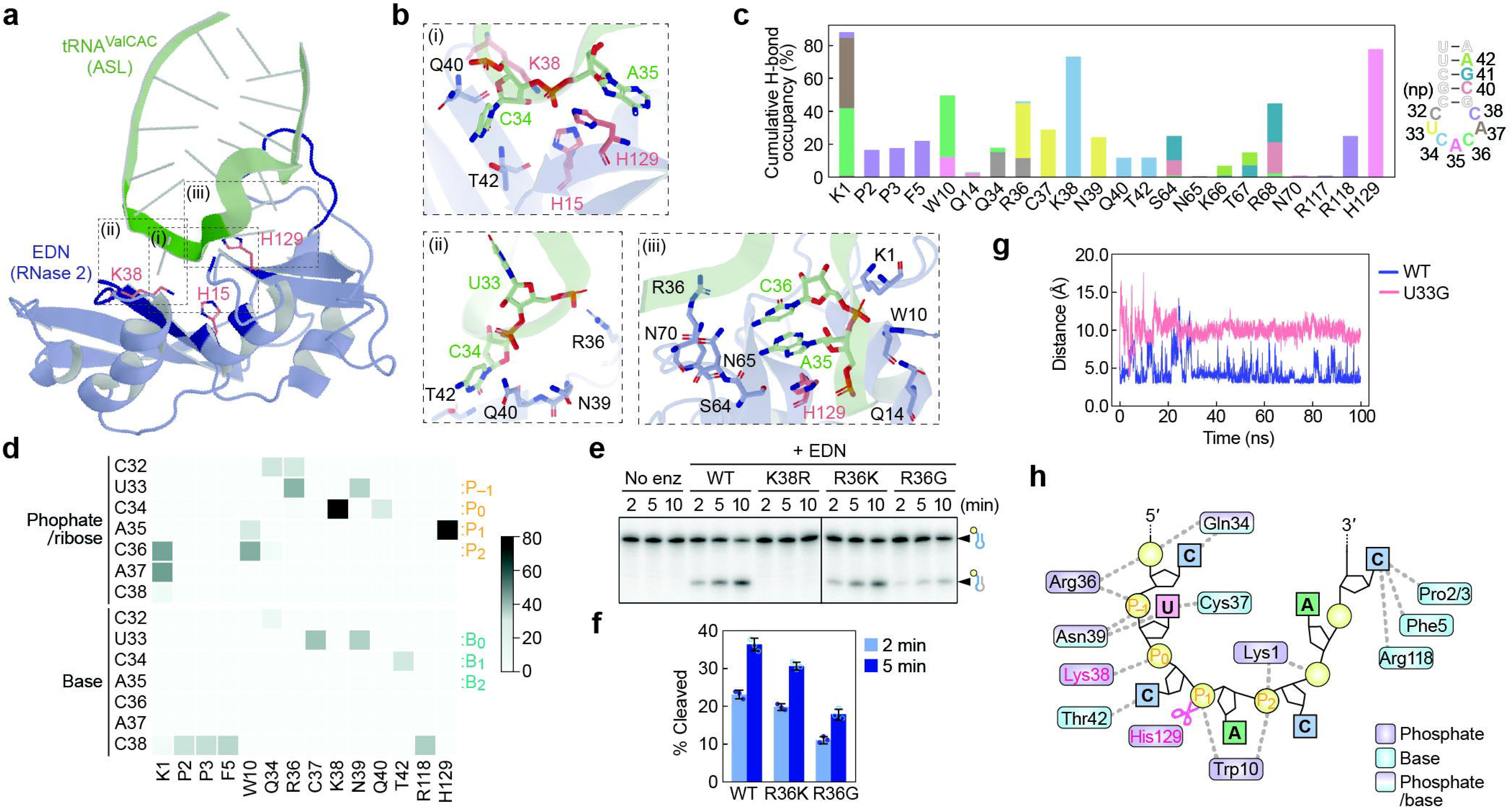
Structural basis of EDN-mediated cleavage of the tRNA anticodon loop. **(a)** Binding model of human EDN with ASL^ValCAC^, derived from the 100-ns endpoint snapshot of an MD simulation. **(b)** Close-up views of the boxed regions (i–iii) in (a), highlighting key interactions within the EDN–ASL complex. **(c)** Cumulative hydrogen-bond occupancy between EDN and ASL^ValCAC^ over 100 ns of MD simulation. **(d)** Hydrogen-bond occupancy at key EDN residues interacting with ASL^ValCAC^; color intensity reflects bond occupancy. **(e)** *In vitro* cleavage assay of Val^modi^ by WT EDN and indicated mutants. **(f)** Quantification of cleavage efficiency in (e). Error bars, S.D. from three independent experiments. **(g)** Time-dependent distances between Arg36 of EDN and the phosphate at np 33 (P_–1_) in ASL^ValCAC^ and its U33G mutant. **(h)** Schematic representation of EDN recognition of ASL^ValCAC^.

Within the catalytic center, the phosphate group between C34 and A35 (P_1_: numbered according to previous reports ^45,60^), the experimentally validated cleavage site, was coordinated by catalytic residues His15, Lys38, and His29 (**Fig. 5b**, top left, **and Extended Data Fig. 6c**). Consistent with prior evidence that Thr42, a residue conserved across the RNase A superfamily, governs cytosine specificity at the cleavage site ^61^, its side-chain hydroxyl group formed a stable hydrogen bond with the N3 of the C34 base (**Fig. 5b**, bottom left). Additional interactions were observed between the 3′-region of the ASL and multiple residues, including Lys1, Trp10, and loop 5 residues (Ser64–Arg68), particularly engaging anticodon bases A35 and C36 (**Fig. 5b**, bottom right), thereby reinforcing the correct alignment of the loop within the catalytic cleft. Importantly, Arg118 formed a hydrogen bond with C38, a nucleotide enriched in EDN-favored tRNAs (**Fig. 4d**), while some N-terminal residues (Pro2, Pro3, and Phe5) also contributed to C38 recognition (**Fig. 5c**). These specific interactions were corroborated by residue-level binding free energy calculations, which indicated favorable energetic contributions consistent with a stabilizing role in the EDN–ASL complex (**Supplementary Table S1**).

## EDN anchors phosphate backbones at the 5′-side of the anticodon loop to mediate efficient cleavage

Hydrogen bond occupancy analysis revealed that residues Arg36, Cys37, and Asn39 of EDN engage in stabilizing interactions with the conserved U33 in the anticodon loop (**Fig. 5c**). The heatmap, separately showing hydrogen bonds with the bases and the phosphate-sugar backbone, indicates that Arg36 interacts with the phosphate-sugar backbone of U33 (P_–1_) (**Fig. 5d and Extended Data Fig. 6d**). Similarly, the phosphates 3′-downstream of the scissile site are supported by Lys1 and Trp10. These multiple interactions with the phosphate backbone suggest roles in substrate positioning and electrostatic stabilization. Notably, no significant interaction was observed at A37 or its downstream phosphate, which were slightly distant from EDN, explaining why the A37U ASL mutant did not show reduced cleavage efficiency (**Fig. 4h**). To assess the functional significance of the phosphate interaction mediated by Arg36, we generated two EDN mutants: R36K and R36G. The R36G mutation markedly impaired catalytic activity, whereas the R36K mutant, which retains a positively charged side chain, exhibited activity comparable to the wild-type (**Fig. 5e-f**). These results underscore the importance of the positive charge at this position for phosphate-backbone recognition.

Next we simulated the interaction between EDN and a U33G mutant ASL to investigate the role of conserved U33. Although the global RMSD remained comparable to the wild-type ASL (**Extended Data Fig. 6e**), the distance between Arg36 and the phosphate group of G33 (P_–1_) increased substantially (**Fig. 5g**), and the hydrogen bond occupancy with those anchoring the phosphate backbone, Gln34 and Arg36, decreased (**Extended Data Fig. 6f**). Consistent with the unchanged *K*_m_ of the U33G mutant (**Fig. 4j**), these results suggest that EDN does not critically recognize the base identity at position 33 but instead utilizes phosphate-backbone anchoring to properly orient the scissile site (P_1_) within the catalytic center.

Together, these findings establish a mechanistic model in which EDN recognizes structural features of the tRNA anticodon loop by anchoring the phosphate backbone both upstream and downstream of the scissile phosphate (**Fig. 5h**). This engagement explains EDN’s preferential recognition of specific tRNA substrates and its consistent cleavage at or near np 34. The residues mediating this interaction, such as Lys1 and Arg36, are conserved in RNase A, RNase 7, and RNase 8, but not in other RNase A superfamily members (**Extended Data Fig. 6c**). This divergence likely reflects specialized biological roles of EDN, distinct from those of its paralogs, and could underlie its generation of immunomodulatory tRNA halves in asthma.

## Discussion

In this study, we identify EDN as the endoribonuclease that selectively cleaves tRNA anticodon loops and generates immunoactive tRNA halves in asthma. These findings elucidates a long-sought biological role for a member of the RNase A superfamily, linking its enzymatic activity to immune response through site-specific tRNA processing. Although RNase A family enzymes have served as paradigms of acid–base catalysis for over half a century ^62^, their endogenous RNA substrates and pathophysiological functions have remained elusive. Our data demonstrate that EDN, an exclusively upregulated ribonuclease in asthma, preferentially cleaves a defined subset of tRNAs, primarily at C–A or other specific motifs within the anticodon loop, producing 5′-tRNA halves that are packaged and secreted in EVs. Among these, 5′-ValCAC and 5′-HisGUG are potent activator of TLR7, capable of amplifying immune responses through cytokine induction. We postulate that EDN upregulation in asthma contributes to disease pathogenesis by generating immunoactive tRNA halves, which in turn potentiate inflammatory changes in the lung. These findings provide a molecular explanation for the long-recognized elevation of EDN in asthma ^38–41^, positioning EDN at the nexus of airway remodeling and inflammation.

Our studies extensively investigated the molecular features that determine substrate selectivity of EDN. This specificity is governed by both the primary nucleotide sequence and the structural features of tRNAs. Unlike previous biochemical studies that relied on dinucleotides or other artificial constructs, our studies tested EDN’s enzymatic properties on its natural target, the tRNA anticodon loops. EDN anchors the phosphate backbone, positioning the scissile phosphate between the first and second anticodon nucleotides within its catalytic pocket. This interaction is mediated by conserved residues: Arg36 and Asn39 anchor the upstream phosphates, while Lys1 and Trp10 stabilize the downstream phosphates. Moreover, conserved nucleotides in tRNAs such as U33 and C38 contribute to EDN recognition. Together, these interactions give rise to asymmetric cleavage at the 5′-side of the anticodon loop in select tRNA species, providing the structural and biochemical basis for the cleavage of endogenous RNA targets by a member of the RNase A superfamily. The residues Arg36 and Asn39 have been suggested to target the anticodon cleavage site of tRNA^AspGUC^ ^45,63^, highlighting their conserved role in substrate recognition.

Our work reframes the RNase A superfamily as a regulated effector arm of airway immunity, capable of reprogramming the RNA landscape to elicit specific receptor responses. The RNase A superfamily has often been referred to as “defensive ribonucleases” due to evidence of their antiviral and antibacterial activities ^64,65^. However, whether such functions operate under physiological conditions has remained unclear. EDN emerges here as a paradigm for this concept, integrating eosinophil activation, site-selective tRNA cleavage, and TLR7 engagement to drive airway inflammation. Notably, TLRs have been postulated to play dual roles, either promoting or suppressing airway inflammation depending on context ^66–69^. Given that eosinophil activation is one of the most prominent immune responses upon antigen sensing in the airways ^70^, EDN and subsequently generated immunoactive tRNA halves likely play a crucial role in the initial phases of asthma, linking eosinophil activity to the initiation and propagation of asthma pathology and serving as key mediators of the epithelial–immune crosstalk. In summary, we propose that EDN and tRNA halves contribute to asthma pathogenesis, with tRNA halves potentially serving as diagnostic biomarkers and EDN-mediated production of tRNA halves representing a prospective therapeutic target.

## Methods

### Ethics statement

All mouse experiments were conducted in compliance with NIH guidelines and approved by the Institutional Animal Care and Use Committee at Thomas Jefferson University (TJU).

### Mouse models of allergic asthma

Intranasal challenges with HDM were performed as described previously ^71^. Briefly, 8-week-old female C57BL/6 mice received 25 μg of HDM extract (*Dermatophagoides pteronyssinus*, Greer Labs) in 35 μl saline intranasally, 5 days per week for 3 consecutive weeks. Lungs were harvested for analysis 24 h after the final challenge.

### Human cell lines and EDN treatment

HeLa (CCL-2), A549 (CCL-185), and BEAS-2B (CRL-3588) cells were obtained from ATCC. HeLa and A549 cells were cultured in DMEM with 10% FBS, and BEAS-2B cells were maintained in DMEM/F12 with 10% FBS. Cells were treated with recombinant EDN at 2 μg/ml and harvested by trypsinization to remove residual EDN bound to the cell surface.

### RNA isolation and northern blot

Total RNA was extracted from frozen tissues and cultured cells using TRIsure (Bioline), following the manufacturer’s instructions for cells and a published protocol for tissues ^26^. Northern blot was performed as previously described ^6,16,27^ using the following antisense probes: 5′-ValCAC, 5′-GAGGCGAACGTGATAACCACTA-3′; 5′-HisGUG, 5′-GCAGAGTACTAACCACTATACG-3′; 5′-iMetCAU, 5′-GCTTCCGCTGCGCCACTCT-3′; 5′-LysCUU, 5′-GTCTCATGCTCTACCGACTG-3′; and 5S rRNA, 5′-GTTCAGGGTGGTATGGCCGT-3′.

### RNA-seq and bioinformatics

For mouse lungs, sncRNAs (20–50 nt) were gel-purified from the total RNA and subjected to cP-RNA-seq procedure using TruSeq Small RNA-seq Library Preparation Kits (illumina) as previously described ^6,14,16,24,26,27,72^. Amplified cDNAs were gel-purified and sequenced on Illumina NextSeq 500 at the MetaOmics Core Facility of the Sidney Kimmel Comprehensive Cancer Center (SKCCC) at TJU. The cP-RNA-seq libraries generated approximately 24–28 million raw reads. For BEAS-2B cells, sncRNAs were isolated using the mirVana miRNA Isolation Kit (Thermo Fisher Scientific) and sequenced by cP-RNA-seq, producing ∼2–10 million raw reads. The sequence data are available in the NCBI Sequence Read Archive under accessions PRJNA525110 and PRJNA1099176.

Bioinformatics analyses followed published pipelines ^26,27^. Briefly, reads were quality filtered and those between 20–50-nt (mouse lungs) and 15–55-nt (BEAS-2B) were extracted using Cutadapt (doi.org/10.14806/ej.17.1.200). Filtered reads were mapped to cytoplasmic mature tRNAs, rRNAs, mRNAs, mitochondrial transcripts, and the whole genome using Bowtie2. The reference tRNA sequences were obtained from Genomic tRNA database (Release 18)^73^. The other reference sequences were derived from GRCm38/mm10 and GRCh37/hg19. Expression and sequence analyses, as well as data visualization, were performed using FastQC and several R packages, including DESeq2 ^74^, *ggplot2*, and *gplots*.

For the analysis of mRNA expression in mouse lungs, we selected four independent projects: PRJNA212775 ^30^, PRJNA547063 ^31^, PRJNA522698, and (iv) PRJNA480707 ^32^. The STAR aligner ^75^ was employed for mapping. The phylogenetic tree was created using MEGA11 ^76^. For the analysis of mRNA expression in human blood, we used the PRJNA415959 dataset and mapped to human transcripts using Bowtie2.

### Quantification of RNAs by qPCR

Specific species of 5′-tRNA halves were quantified using TaqMan-based RT-qPCR as described previously ^6,14,16,26^. Briefly, total RNA was treated with T4 polynucleotide kinase (T4 PNK, New England Biolabs) to remove cP from the 3′-end of 5′-tRNA halves, followed by ligation to a 3′-RNA adaptor using T4 RNA ligase (Thermo Fisher Scientific). The ligated RNA was then subjected to RT-qPCR using the One Step PrimeScript RT-PCR Kit (Takara Bio), with a TaqMan probe targeting the boundary of the target RNA and the 3′-adaptor, along with forward and reverse primers. ActB mRNA was quantified by standard RT-qPCR as described previously. Sequences of synthetic oligonucleotides used for qPCR are available in previous reports. The information on license plates ^77^ and names via tDRnamer ^78^ for the targeted 5′-tRNA halves are included in **Supplementary Tables S2**.

### Purification of recombinant EDN

The coding region of the human *RNase 2 (EDN)* gene was cloned into the pET-21a plasmid to express C-terminal 6× His-tagged EDN protein. K38R, R36G, and R36K mutant plasmids were produced by site-directed mutagenesis. The wild-type and mutant EDN were expressed in *E. coli* BL21 (DE3) pLysS as described previously ^79^. Cells were lysed by sonication in a lysis buffer (50 mM Tris-HCl pH8.0, 2 mM EDTA, 0.5% Triton-X100, 10% glycerol, and 0.7 mg/ml lysozyme). The insoluble fraction, containing recombinant EDN, was recovered by centrifugation, denatured, and refolded in the presence of oxidized glutathione as described previously ^80^. Refolded protein was dialyzed overnight against IEX buffer A (20 mM Tris-HCl pH7.5, 0.2% Triton X-100, and 2% glycerol), centrifuged at 20,000 ×*g* for 20 min, and filtered through a 0.45 μm membrane. The filtered solution was applied to a HiTrap SP HP column (Cytiva) and eluted with IEX buffer B (20 mM Tris-HCl pH7.5, 1 M NaCl, 0.2% Triton X-100, and 2% glycerol). EDN-containing fractions were pooled, concentrated on a Vivaspin 500, 5k MWCO. The purified protein was stored in a storage buffer containing 10 mM Tris-HCl pH7.5, 50 mM NaCl, 2 mM DTT, and 40% glycerol. Ribonucleolytic activity of purified EDN was assessed using the Kunitz assay that measures the release of nucleotides into the acid soluble fraction as previously described ^81^.

### Western blot

Western blot was performed as described previously ^6,16^. Anti-His6 monoclonal antibody (6*His, Proteintech), anti-GAPDH monoclonal antibody (Proteintech), anti-Alix (Proteintech), and anti-Cytochrome *c* (Santa Cruz Biotechnology) were used as the primary antibodies.

### EV isolation

EVs were isolated from the culture medium of A549 cells using an ultracentrifugation-based method, as described previously ^6,7^. Successful EV isolation was verified by western blot and NTA as per our previous study ^6^.

### *In vitro* RNA synthesis

Synthetic tRNA^ValCAC^ and tRNA^HisGUG^ were produced by *in vitro* transcription as described previously ^6,16,82^. Briefly, dsDNA templates were generated using T7 DNA Polymerase (New England Biolabs) and transcribed with T7 RNA polymerase (New England Biolabs) at 37°C for 6 h. The synthesized RNAs were gel-purified using denaturing PAGE with single-nucleotide resolution. DNA/RNA chimeric ASLs and RNA ASLs were synthesized by Integrated DNA Technologies. The sequences of the synthetic RNAs are shown in **Supplementary Table S3**.

### *In vitro* RNA cleavage assay

Substrate RNAs or DNA/RNA chimeric constructs were 5′-end labeled with γ-^32^P-ATP using T4 PNK and purified with Centri-Spin 10 columns (Princeton Separations). Trace amounts of labeled oligos were mixed with the indicated concentration of unlabeled (cold) constructs and incubated with recombinant EDN at 37°C for the indicated time. Unless otherwise specified, reactions were performed in a buffer containing 50 mM MES pH 6.0, 25 mM NaCl, and 0.1 mg/mL BSA, and MgCl_2_ (20 mM for tRNAs or 10 mM for ASLs). Cleavage products were resolved by denaturing PAGE (12% acrylamide/bis-acrylamide for tRNAs or 20% for ASLs) and visualized on Typhoon phosphorimager (Cytiva). Signal intensities were quantified using ImageJ ^83^. For native HeLa tRNA fractions, 500 ng of gel-purified tRNAs were incubated with 0, 1, 5, or 25 nM of EDN for 1 min in a buffer containing 50 mM MES pH5.5, 20 mM MgCl_2_, and 0.1 mg/ml BSA, followed by northern blot. The cleavage assay for Michaelis–Menten kinetics was performed using substrate concentrations of 0.2–18 μM. Initial velocities were determined from the linear portion of the reaction progress curves and fitted to the standard Michaelis–Menten equation using GraphPad Prism.

### Molecular Docking and MD simulations

The initial EDN–ASL complex model was constructed by extracting the ASL^ValCAC^ from the *Thermus themophilus* tRNA^ValCAC^–valyl-tRNA synthetase crystal structure (PDB 1IVS) ^57^. The ASL was positioned by superimposing the active site of EDN (PDB 1HI3) ^84^ onto that of RNase A in complex with d(CpA) (PDB 1RPG) ^58^, with the C34–A35 dinucleotide of the ASL aligned to the CpA substrate. The U33G mutant structure was generated by substituting U33 with G in the wild-type ASL model using the mutagenesis function in PyMOL (The PyMOL Molecular Graphics System, Version 3.1.0 Schrödinger, LLC). MD simulations were performed using GROMACS 2023.2 ^85^ with the AMBER99SB-ILDN force field ^86^. Each complex was solvated using the SPC216 water model in a cubic box and neutralized with 150 mM NaCl. Electrostatic and van der Waals interactions were truncated at 1.0 nm, and long-range electrostatics were calculated with the particle mesh Ewald method ^87^. Energy minimization was performed using the steepest descent algorithm, and the system was then equilibrated at 300 K and 1 bar for 100 ps each in two steps: first under constant volume (NVT), then under constant pressure (NPT). Temperature and pressure were controlled using the V-rescale thermostat and Parrinello-Rahman barostat, respectively ^88^. The LINCS algorithm ^89^ was used to constrain bonds involving hydrogen atoms. The integration time step was set as 2 fs. Production runs of 100 ns were conducted under the NPT ensemble without restraints, with three independent replicates for each complex (wild-type and U33G mutant). The RMSD and hydrogen bonds profiles were generated by GROMACS modules. The total binding free energy, van der Waals energy and electrostatic energy of the 100 ns production runs were calculated by using the gmx_MMPBSA^90^.

Structural prediction using AlphaFold3 was performed as follows: the input consisted of the EDN amino acid sequence without the signal sequence and a 15-nt RNA fragment corresponding to the ASL of tRNA^ValCAC^. The top-ranked prediction model was used for visualization, and its reliability was evaluated through the predicted local distance difference test (pLDDT) value and the predicted aligned error (PAE) matrix generated by alphafold3_tools (https://github.com/cddlab/alphafold3_tools). Images were generated by PyMOL (The PyMOL Molecular Graphics System, Version 3.1.0 Schrödinger, LLC) and Matplotlib (Version 3.4.3) library of Python (Version 3.10.12).

## Acknowledgements

We are grateful to the members of our laboratories for helpful discussions, to Dr. Hideharu Hashimoto for guidance on the fundamentals of kinetic analysis, and to Abigail Sims for technical assistance with bacterial culture. This study was supported by the National Institutes of Health Grant (AI130496 and HL175371 to D.A.D and Y.K., GM106047, GM152185, and AI171366 to Y.K., and HL137030 to D.A.D), the American Cancer Society Research Scholar Grant (RSG-17-059-01-RMC, to Y.K.), the W.W. Smith Charitable Trust Grant (C1608, to Y.K.), the Japan Society for the Promotion of Science Postdoctoral Fellowship for Research Abroad (to M.S.), the American Lung Association Catalyst Award Grant (to M.S.), and the Japan Science and Technology Agency Support for Pioneering Research Initiated by the Next Generation (JPMJSP2108 to A.M.). The authors declare no competing financial interests.

## Contributions

D.A.D. and Y.K. conceived and initiated the study, and M.S. conceived the subsequent development of the study. M.S. and Y.K. supervised the project. M.S. conducted most of the experiments, except for mouse HDM treatment, which was carried out by S.D.S. under the supervision of D.A.D., and EV isolation and its analyses, which were performed by T.K. M.S. performed the bioinformatic analysis of RNA-seq data. M.S. also performed the kinetic assay with input from O.T. A.M. performed the MD simulations with input from M.S. and O.T. M.S. and Y.K. drafted the original manuscript. All authors reviewed the manuscript and approved the final version for submission.

