## Supplemental Information for "RNase 2/EDN cleaves tRNA anticodon loops to generate immunoactive RNAs"

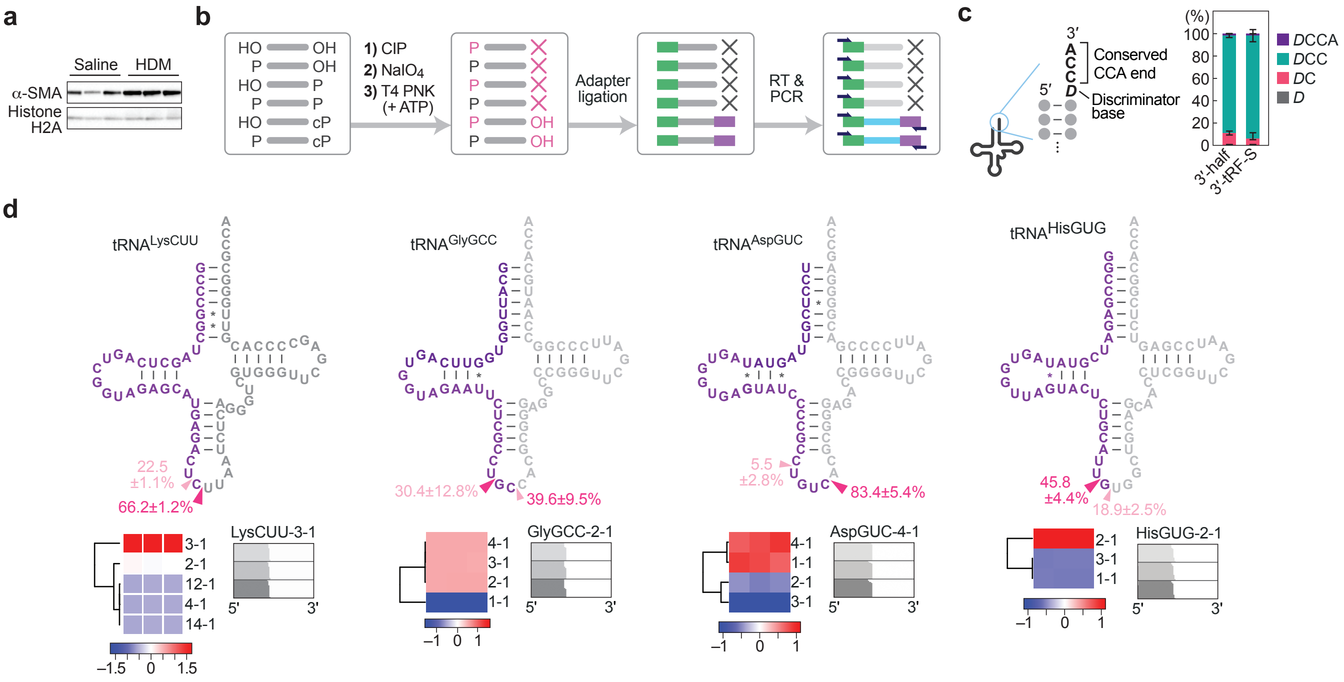

**Extended Data Figure 1.**
**(a)** Western blot of mouse lung lysates showing upregulation of α-SMA following HDM challenge. **(b)** Schematic of the cP-RNA-seq workflow. sncRNAs are sequentially treated with calf intestinal alkaline phosphatase (CIP) and sodium periodate ($NaIO_4$), leaving only the cP end intact. The cP is then converted to a 3′-OH by T4 PNK treatment, enabling 3′-adaptor ligation, and allowing selective amplification and sequencing of sncRNAs originally bearing a cP. **(c)** Variation at the 3′-end of 3′-tRNA halves and 3′-tRFs-S. **(d)** Cleavage patterns for 5′-tRNA halves from selected isoacceptors. Cloverleaf structures indicate major cleavage sites (arrowheads). Errors represent the S.D. from three replicates. Heatmaps show relative abundance of isodecoders; most abundant isodecoder alignments are also shown.

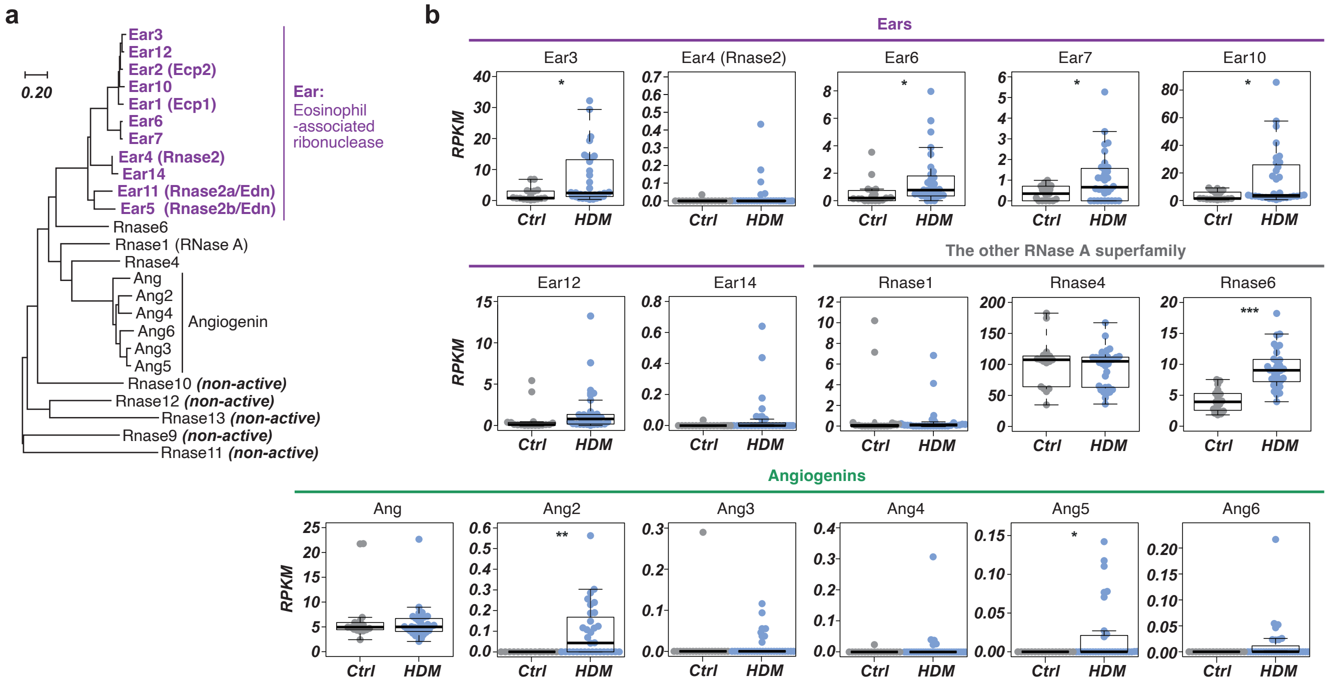


**Extended Data Figure 2.**

**(a)** Phylogenetic tree of murine RNase A superfamily members.

**(b)** mRNA expression levels of other RNase A superfamily members in mouse lungs (the datasets analyzed in Fig. 2a). Student's *t*-test: *$p$<0.05; **$p$<0.01; ***$p$<0.001.

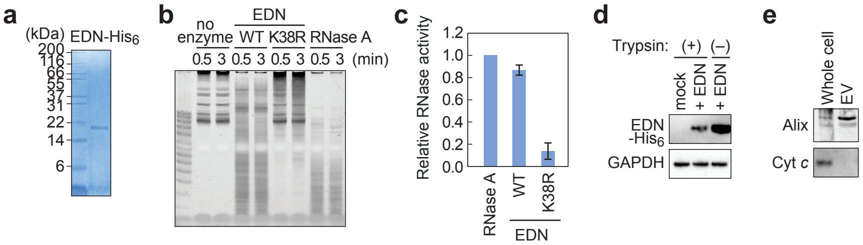


**Extended Data Figure 3.**
**(a)** Recombinant human EDN analyzed by SDS-PAGE and stained by CBB. **(b)** Ribonuclease activity of EDN assessed by incubating with HeLa total RNA. Cleaved RNAs were stained with SYBR Gold. **(c)** Quantification of EDN activity by Kunitz assay. Error bars indicate S.D. obtained from three independent experiments. **(d)** Western blot showing recombinant EDN in HeLa cells after 4 h of treatment, with or without trypsinization prior to harvest. Cells harvested without trypsinization exhibited higher EDN levels when equal amounts of total protein were loaded, indicating that the elevated signal reflects surface-bound EDN rather than internalized protein. **(e)** Western blot of cell extract and EV fraction obtained from A549 cells. The EV fraction contained the EV marker Alix but not the non-EV protein Cyt *c*.

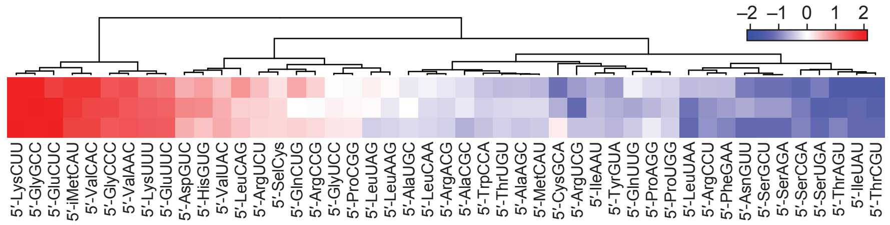


**Extended Data Figure 4.**

Abundance of 5′-tRNA halves in EDN-treated BEAS-2B cells. Color bar indicates *z*-scores.

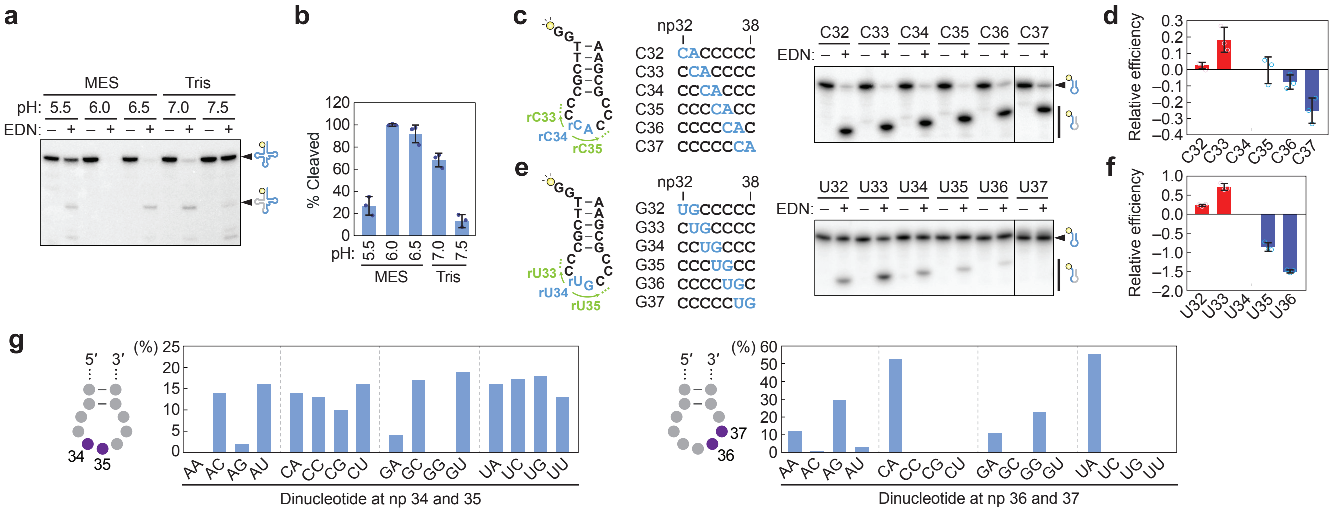

**Extended Data Figure 5.**

**(a)** pH dependence of EDN activity. 5′-$^{32}$P-labeled tRNAValCAC was incubated with EDN at the indicated pH. **(b)** Cleavage efficiency in (a) was quantified from three independent experiments. Error bars, S.D. from three independent experiments. **(c, e)** Cleavage assay using DNA-RNA chimeric ASL$^{ValCAC}$ (c) or ASL$^{HisGUG}$ (e) variants, in which the rC–A or rU–G cleavage site was systematically shifted from np 32 to 37. **(d, f)** Relative cleavage efficiencies of ASL variants in (c) and (e). Cleavage of the C34 (ASL$^{ValCAC}$) or U34 (ASL$^{HisGUG}$) construct was set to 1, and fold changes for other substrates are presented on a $\log_2$ scale. Error bars represent S.D. from three independent experiments. **(g)** Frequency of dinucleotides at the indicated positions within ASLs of human tRNA genes.

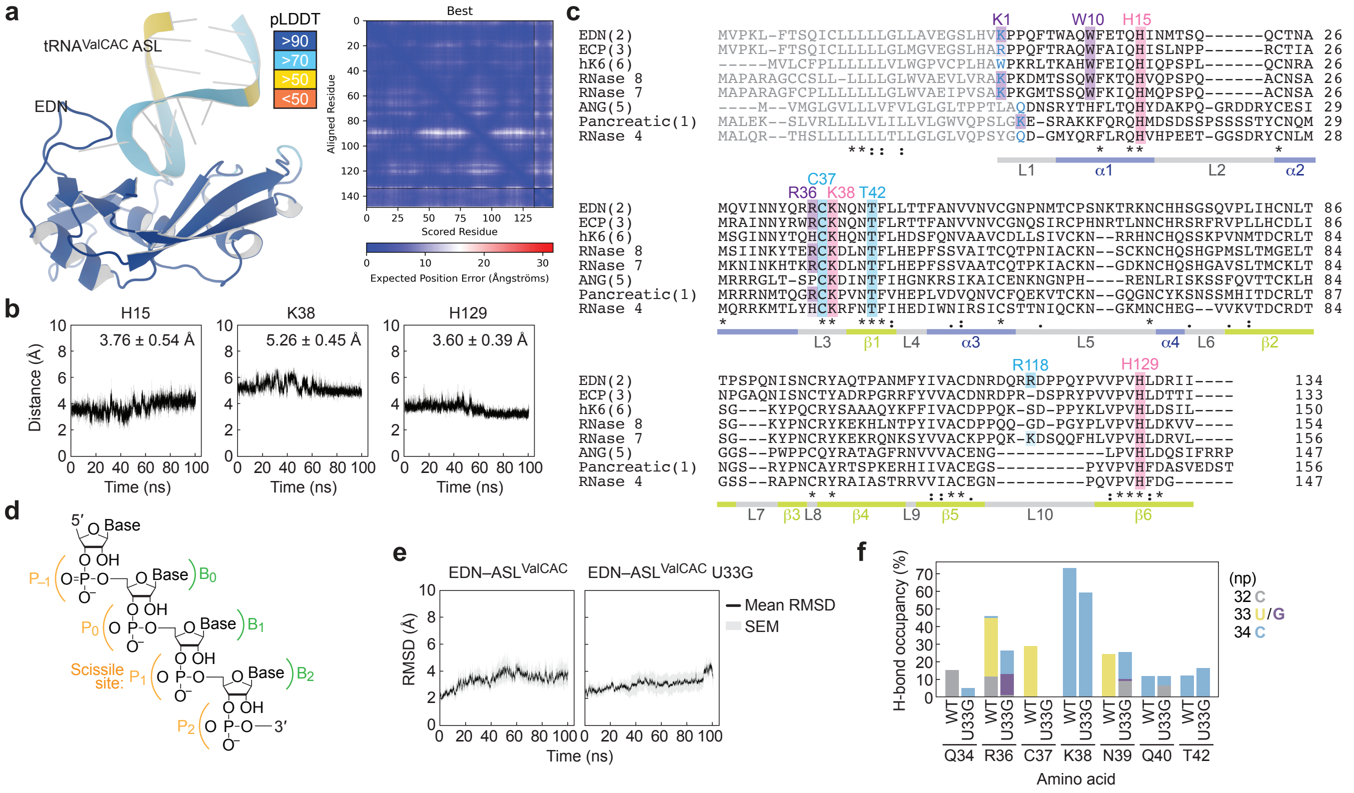

**Extended Data Figure 6.**

**(a)** Structure model of the EDN–ASL$^{ValCAC}$ complex predicted by AlphaFold3. Shown are a ribbon representation of the complex colored by pLDDT (left) and the PAE matrix (right). **(b)** Time-dependent distances between the three catalytic residues of EDN (H15, K38, H129) and the cleavage site nucleotide, monitored over 100 ns of MD simulation. Mean ± S.D. is indicated. **(c)** Sequence alignment of RNase A superfamily members highlighting the conserved catalytic residues (H15, K38, H129) and substrate recognition residues. **(d)** Numbering scheme of phosphates and bases surrounding the scissile site. **(e)** RMSD profiles of the EDN–ASL$^{ValCAC}$ and EDN–ASL$^{ValCAC}$ U33G complexes over 100 ns MD simulations. **(f)** Hydrogen bond occupancy of selected EDN residues interacting with C32, U/G33, and C34 in ASL$^{ValCAC}$ or its U33G mutant.

**Table S1. Average binding free energy of each replicate (kcal/mol)**

| EDN | Replicate | ΔG_total | ΔVDW | ΔEEL | ΔEPB |
|---|---|---|---|---|---|
| WT | 1 | -64.76 ± 2.31 | -71.01 ± 0.89 | -2176.39 ± 16.72 | 2190.13 ± 16.50 |
| | 2 | -77.12 ± 3.71 | -78.71 ± 1.11 | -2222.02 ± 14.91 | 2231.78 ± 14.24 |
| | 3 | -55.16 ± 1.95 | -72.93 ± 1.54 | -2139.75 ± 18.32 | 2165.30 ± 18.66 |
| U33G | 1 | -54.93 ± 2.86 | -67.58 ± 1.31 | -2131.63 ± 14.60 | 2151.37 ± 15.29 |
| | 2 | -66.08 ± 3.60 | -87.97 ± 2.97 | -2244.22 ± 18.71 | 2274.46 ± 18.87 |
| | 3 | -56.90 ± 2.10 | -65.68 ± 1.11 | -2159.77 ± 18.05 | 2175.55 ± 17.94 |

**Table S2. Unique ID for 5′-halves experimentally investigated in this study**

| Name | Sequence (5′-to-3′) | License plate* | tDRname** |
|---|---|---|---|
| 5′-GluCUC | UCCCUGGUGGUCUAGUGGUUAGGAUUCGGCGCUC | tRF-34-87R8WP9N1EWJI5 | tDR-1:34-Glu-GTC-1 |
| 5′-iMetCAU | AGCAGAGUGGCGCAGCGGAAGCGUGCUGGGCCC | tRF-33-FP18LPMBQ4NKDJ | tDR-1:33-Met-CAU-1 |
| 5′-ValCAC | GUUUCCGUAGUGUAGUGGUUAUCACGUUCGCCUC | tRF-34-79MP9P9NH57S15 | tDR-1:34-Val-CAC-1-M3 |
| 5′-HisGUG | GCCGUGAUCGUAUAGUGGUUAGUACUCUGCGUU | tRF-33-PW5SVP9N15WV0E | tDR-1:33-His-GTG-1 |

*Pliatsika, V., et al. *Bioinformatics* (2016)

*Holmes, A.D., et al. *Nature Methods* (2023)

**Supplementary Table S3. Sequences of synthetic oligonucleotides used for biochemical assays**

| ID | Variant ID | Sequence (5′-to-3′) |
|---|---|---|
| Mature tRNAValCAC | N/A | GUUUCCGUAGUGUAGUGGUUAUCACGUUCGCCUCACACGCGAAAGGUCCCCGGUUCGAAACCGGGCGGAAACACCA |
| Mature tRNAHisGUG | N/A | GGCCGUGAUCGUAUAGUGGUUAGUACUCUGCGUUGUGGCCGCAGCAACCUCGGUUCGAAUCCGAGUCACGGCA |
| Chimeric ValCAC-ASL | rCA | GGTTCGCCTrCACACGCGAA |
| | rCT | GGTTCGCCTrCTCACGCGAA |
| | rCG | GGTTCGCCTrCGCACGCGAA |
| | rCC | GGTTCGCCTrCCCACGCGAA |
| | rUA | GGTTCGCCTrUACACGCGAA |
| | rUT | GGTTCGCCTrUTCACGCGAA |
| | rUG | GGTTCGCCTrUGCACGCGAA |
| | rUC | GGTTCGCCTrUCCACGCGAA |
| | rGA | GGTTCGCCTrGACACGCGAA |
| | rGT | GGTTCGCCTrGTCACGCGAA |
| | rGC | GGTTCGCCTrGCCACGCGAA |
| | rAA | GGTTCGCCTrAACACGCGAA |
| | C32 | GGTTCGCrCACCCCCGCGAA |
| | C33 | GGTTCGCCrCACCCCGCGAA |
| | C34 | GGTTCGCCCrCACCCGCGAA |
| | C35 | GGTTCGCCCCrCACCGCGAA |
| | C36 | GGTTCGCCCCCrCACGCGAA |
| | C37 | GGTTCGCCCCCCrCAGCGAA |
| Chimeric HisGUG-ASL | G32 | GGTTCGCrUGCCCCCGCGAA |
| | G33 | GGTTCGCCrUGCCCCGCGAA |
| | G34 | GGTTCGCCCrUGCCCGCGAA |
| | G35 | GGTTCGCCCCrUGCCGCGAA |
| | G36 | GGTTCGCCCCCrUGCGCGAA |
| | G37 | GGTTCGCCCCCCrUGGCGAA |
| ValCAC-ASL | Valmodi | GUUCGGCUCAGACCCGAA |
| | U33G | GUUCGGCGCAGACCCGAA |
| | A37U | GUUCGGCUCAGUCCCGAA |
| | C38A | GUUCGGCUCAGAACCGAA |
| | U33G/C38A | GUUCGGCGCAGAACCGAA |
| iMetCAU-ASL | iMetmodi | GGCUGGGCCCAGAACCCAG |
